# Exploring *Cryptococcus neoformans* capsule structure and assembly with a hydroxylamine-armed fluorescent probe

**DOI:** 10.1101/871665

**Authors:** Conor J. Crawford, Radamés J.B. Cordero, Lorenzo Guazzelli, Maggie P. Wear, Anthony Bowen, Stefan Oscarson, Arturo Casadevall

**Author notes:** Department of Pharmacy, Università di Pisa, Via Bonanno 6, 56126, Pisa, Italy (Lorenzo Guazzelli). SO and AC share senior authorship. Correspondence can be addressed to either: Stefan Oscarson, Centre for Synthesis and Chemical Biology, University College Dublin, Belfield, Dublin 4, Ireland or Arturo Casadevall, Dept. of Microbiology and Molecular Immunology, Bloomberg School of Public Health, The Johns Hopkins University, 615 N. Wolfe St., Rm. E5132, Baltimore, MD 21205.

## Abstract

Chemical biology is an emerging field that allows the study and manipulation of biological systems using probes that inform on structure based on their reactivity. We report the synthesis of a hydroxylamine-armed fluorescent probe that reacts with reducing glycans and its application to study the architecture of the *Cryptococcus neoformans* capsule under a variety of conditions. The probe signal localized intracellularly and at the cell wall-membrane interface, implying the presence of reducing end glycans at this location where the capsule attachment to the cell body occurs. In contrast, there was no fluorescence signal in the body of the capsule. We observed vesicle-like structures containing the reducing-end probe, both intra- and extracellularly, consistent with the importance of vesicles in capsular assembly. Disrupting the capsule with DMSO, ultrasound, or mechanical shear-stress resulted in capsule alterations that affected the binding of the probe as reducing ends were exposed, and cell membrane integrity was compromised. In contrast to the polysaccharides in the assembled capsule, isolated exopolysaccharides contained reducing ends. The reactivity of the hydroxylamine-armed fluorescent probe suggests a model for capsule assembly where reducing ends localize to the cell wall surface, supporting previous work suggesting that this is an initiation point for capsular assembly. Chemical biology is a promising approach for studying the *C. neoformans* capsule and its associated polysaccharides.

---

*Cryptococcus neoformans* is a major human pathogen causing 1 million infections annually and approximately 600,000 deaths (1). Infection often occurs in early childhood and is followed by either clearance or a state of latency (2, 3). However, if the host immune system becomes compromised through senescence, HIV/AIDS, or is chemically induced for solid organ transplantation, the infection can re-emerge and cause lethal Cryptococcosis or fungal meningitis (1, 4). A further problem is that current anti-fungal drugs often fail to clear an infection in hosts with impaired immunity. A better understanding of the biology of *C. neoformans* may allow the generation of new therapies to combat this infectious disease.

A major virulence factor of *C. neoformans* is the polysaccharide capsule that surrounds the cell body. Polysaccharides are also secreted into the environment and host tissues during infection, known as exopolysaccharide. The Cryptococcal capsule is comprised of a variety of constituents, including: galactoxylomannan (GalXM), mannoproteins, β-glucans, and glucuronoxylomannan (GXM) (5). GXM predominates and contributes to virulence by interfering with host immunity and protecting the fungal cells. The role of these polysaccharides during infection has been implicated in interference and prevention of phagocytosis, inhibition of leukocyte migration, and cytokine production (6–8). The cell wall of *C. neoformans* is also made up of several polysaccharides including β-(1→3) and β-(1→6) glucan, α-(1→3) glucan, chitin, and chitosan (9, 10).

The study of the capsule and its shed exopolysaccharides is challenging because apart from antibodies, we lack tools to probe the fundamental nature of the organization, architecture and structure of the capsule (11). The capsule is composed predominantly of water, which makes it vulnerable to desiccation. Also, little is known regarding how the capsule is synthesized, transported and assembled extracellularly. NMR analysis completed by Cherniak and collaborators defined the chemical structure of the GXM as a linear backbone of α-(1→3)-mannose bearing β-(1→2) and β-(1→4) xylose branches and β-(1→2) glucuronic acid branches, and a 6-*O*-acetyl substitution along the mannan backbone (12). It has been proposed that the glucuronic acid residues mediate divalent cation bridge interactions intramolecularly that help give rise to a polysaccharide architecture through a matrix of self-aggregation (13). Evidence that the capsular polysaccharide consists of a higher-order branched matrix comes from static and dynamic light scattering, viscosity analysis, and high-resolution microscopy (14).

A fundamental question in *C. neoformans* biology is how the capsule is assembled and organized on the cell surface (15). While several glycosidase enzymes required for the biosynthesis of the GXM capsule have been identified by Doering and co-workers (16–18). There is uncertainty over the mechanism of capsule enlargement with two proposed models that are non-mutually exclusive. The first termed ‘proximal growth’ (19), in which the polysaccharide chain is increased in size by incorporation of polysaccharide at the cell body, displacing preexisting molecules to the outer edge (17) i.e. a mechanism consistent with prokaryotic bacteria capsule assembly (20). The second termed ‘distal growth’, where addition of new polysaccharide is incorporated at the capsule edge, with older material remaining close to the cell body (21).

Despite the mechanism of capsule assembly not being elucidated, it is evident that the cell-wall and capsule are capable of dynamic reorganization, as shown during budding events from immunofluorescence studies, such that the mother cell reorganizes its capsule to facilitate the emerging bud, though the exact mechanisms that govern this process remain unclear (11). Several studies suggest intracellular biosynthesis, followed by vesicular transport across the cell wall (22, 23). The importance of vesicles in *C. neoformans* is further demonstrated by mutations in the *C. neoformans* secretory system known as *cap* mutants (22, 24). These mutants, lack capsular GXM but secrete other polysaccharides suggesting the presence of other transport mechanisms independent of that for the capsule, possibly a process that with similarities to that of bacterial capsule biosynthesis (20, 25, 26). It is also unclear how the capsule attachment to the cell wall occurs, or if the cell wall provides the skeleton for proteins or other chemical moieties that then mediate its attachment (9).

Chemical glycobiology is an emerging and exciting sub-discipline of chemical biology that promises to allow further study of the cryptococcal capsule biosynthesis, assembly, and function. Unlike the study of proteins, carbohydrates are not template-encoded, which complicates research in understanding change in their structure and function (27). The introduction of chemical reporter groups on glycans provides new strategies to study an organism’s glycome. Chemical glycobiology has proven the ability to image live cells and organisms using metabolic oligosaccharide engineering (MOE) followed by bio-orthogonal ligation chemistry (28–31). To gain insight into the distribution of reducing sugars in *C. neoformans*, we synthesized a hydroxylamine-armed fluorescent probe (reducing end probe, R.E probe). This structure-activity based probe allowed insight into capsule biosynthesis, structure, and assembly. In a biological setting, hydroxylamine nucleophiles are known to react with a high degree of selectively with the reducing end of glycans, to a give the corresponding oxime conjugate (Figure 1B) (32). This stable adduct, when attached to a fluorophore, would allow imaging of cryptococcal cells and give insight into the distribution of reducing ends in the cell, cell-wall, and capsule.

**Figure 1:**
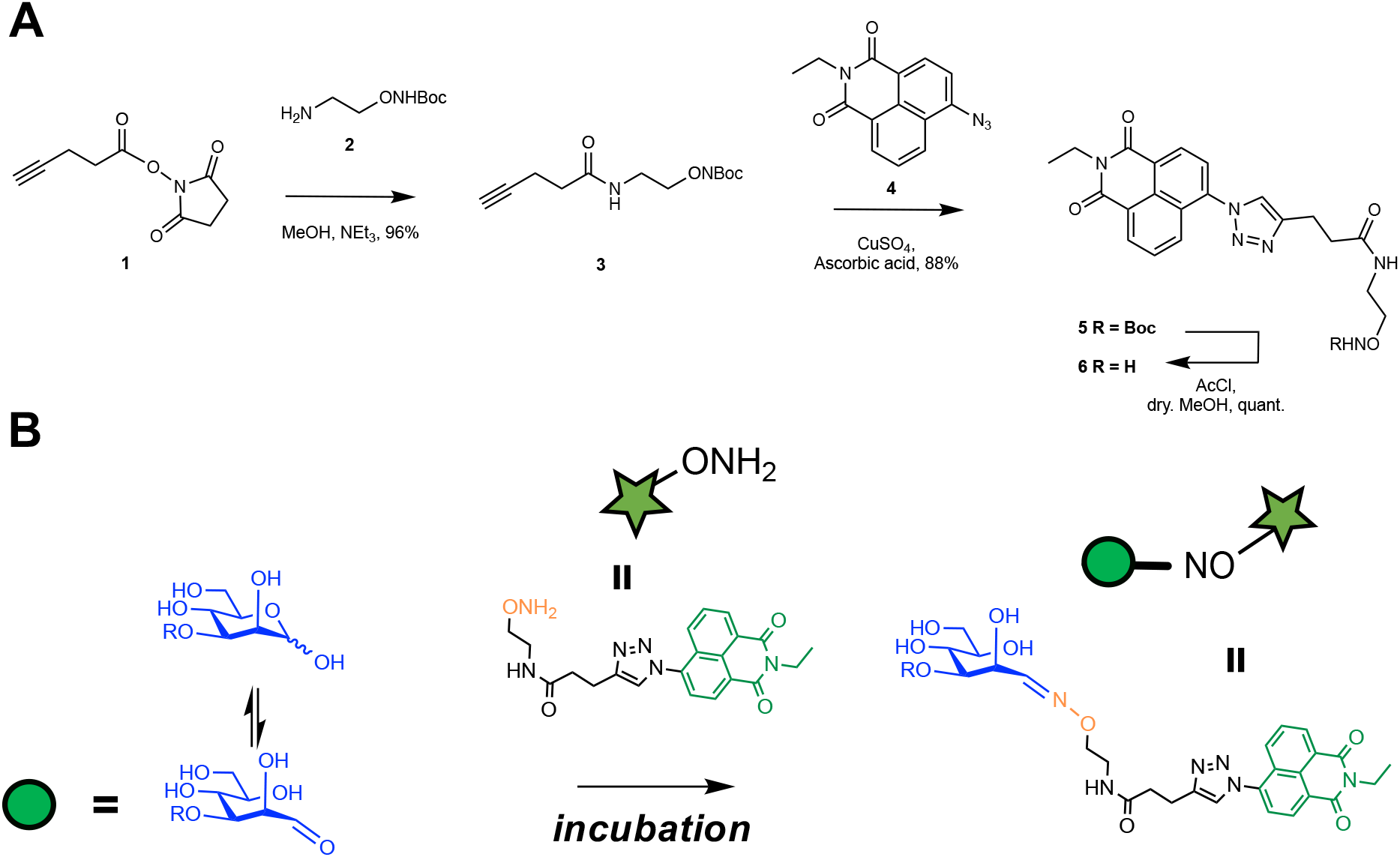
A hydroxylamine-armed fluorescent probe to study the distribution of reducing sugars in *C. neoformans*. **A.** Synthesis of the R.E probe. **B.** Proposed mechanism of action of the R.E armed fluorescent probe. The R.E probe reacts preferentially with aldehydes to form stable oxime adduct which can be used to image the localization of reducing ends in the capsule and inside the cell.

We used this probe to determine the localization of reducing end glycans at the capsule-cell wall interface and visualize the presence of both intra- and extracellular polysaccharide containing vesicles. We also use the probe to examine the effect of different cellular processing techniques on capsular architecture, and that the shed EPS contains reducing ends.

## Results

### Synthesis of hydroxylamine-armed fluorescent probe

To synthesize the hydroxylamine arm of the fluorescent probe, we started from commercially available ethanolamine. After a 5 step synthesis, an *N*-Boc protected hydroxylamine derivative **2** was prepared (33, 34). (Figure 1A) This was then coupled with **1** an *N*-hydroxysuccinimide activated ester of propargylacetic acid (35), after purification by flash column chromatography compound **3** was isolated in a 96% yield. The fluorescent arm of the hydroxylamine-armed probe was prepared from 4-bromo-1,8-naphthalic anhydride, following known literature procedures to give compound **4** (36). The two arms of the probe were then combined using copper click chemistry to produce compound **5** in 88% yield after flash chromatography. Subsequently, the *N*-Boc protecting group was removed using acetyl chloride in anhydrous methanol to produce the hydroxylamine-armed fluorescent probe **6**.

### Incubation of the R.E probe with C. neoformans does not alter fluorescent properties

To confirm that *C. neoformans* was not metabolizing the R.E probe and altering its optical probertites, we incubated the probe with *C*. *neoformans* for 24 hours, washed cells, and performed analysis using a fluorescence spectrometer. Spectrometric analysis confirmed the incorporation of the probe into *C. neoformans* cells (Figure 2A) with characteristic emission maxima of 440-470 nm (excitation 360 nm). The encapsulated H99, *CAP59*Δ and *CAP67*Δ cells displayed higher level of fluorescence compared to that of control cells that were not incubated with the R.E probe (Figure 2A). In contrast, melanized cells displayed no fluorescence emission when excited at this wavelength, likely due to signal quenching of signal by the melanin (37). Encapsulated H99, *CAP59*Δ, and *CAP67*Δ cells also displayed higher RFU at the emission maxima of the probe (Figure 2B).

**Figure 2.**
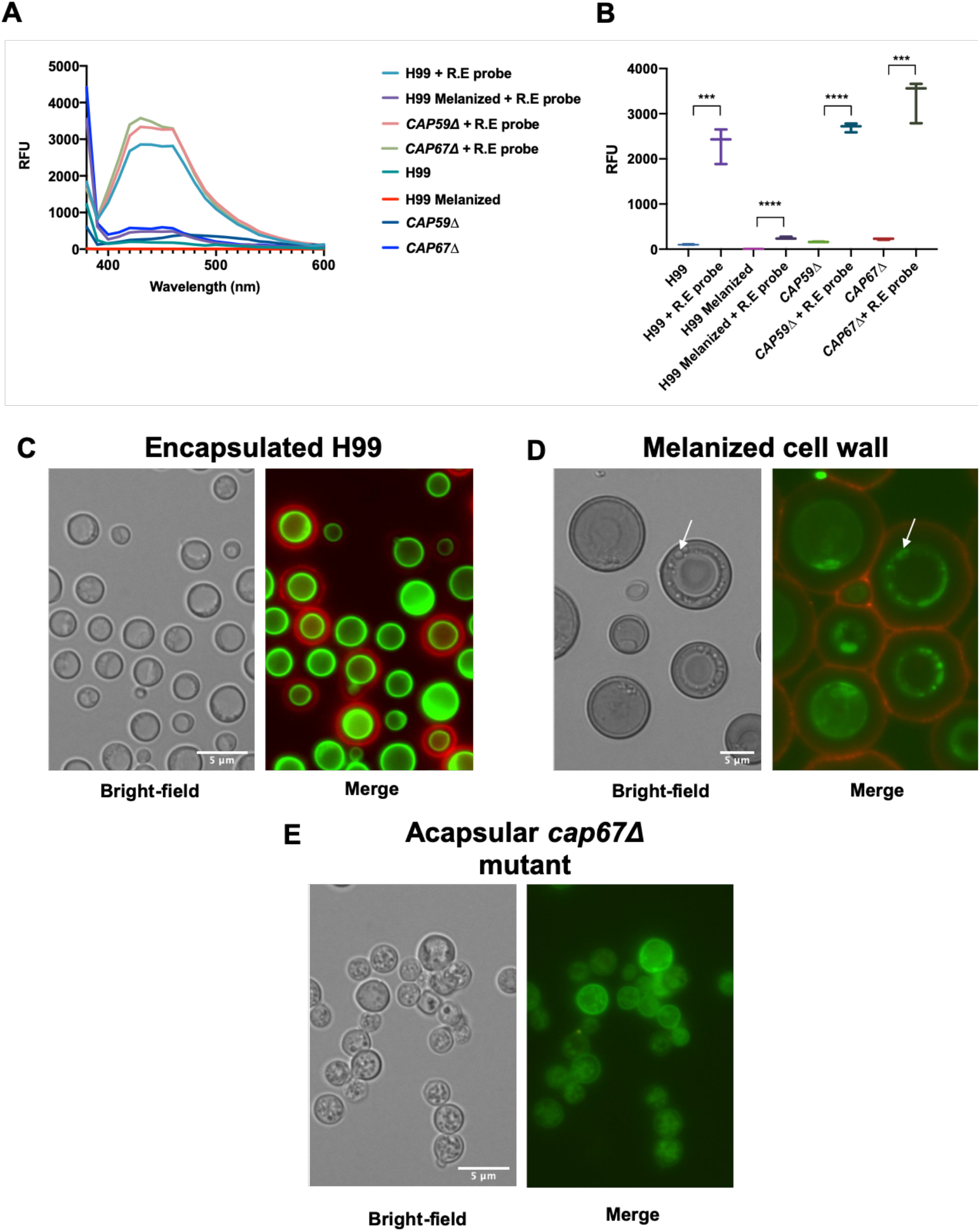
Incubation of hydroxylamine-armed fluorescent probe with *C. neoformans*. **A** Fluorescence spectrum of *C. neoformans* cells incubated with and without hydroxylamine-armed probe (R.E probe), those that have been incubated with the R.E probe have increased emission maxima at ~453 nm (excitation 360 nm) compared to encapsulated H99, *CAP59*Δ, or *CAP67*Δ with the exception of the *C. neoformans* with of encapsulated H99 with a melanized cell wall. Melanization of the cell wall is likely quenching the florescence of the probe. **B** RFU of *C. neoformans* cells incubated with the R.E probe are significantly higher at emission maxima (440 nm) of the probe (excitation 360 nm). Error bars represent 95% confidence interval, Statistical significance was determined using unpaired T-test (*, *p* 0.05; **, *p* 0.01; ****, *p* 0.0001.) **C** Encapsulated H99: The capsule shown by 18B7-AF_594_ (red) displays no reactivity to the R.E probe **6** (green) and is localized at the cell wall-membrane interface. **D** Melanized *C. neoformans* cells display the same localization of the R.E probe. The R.E probe appears intracellularly possibly in vesicular bodies, and displays no reactivity toward capsule, despite the melanization of the cell wall. **E** Acapsular mutant *Cryptococcus neoformans cap67*Δ incubated with the R.E probe shows bright fluorescence intensity coming from the cytoplasm and the cell wall-membrane interface. Labelling of cytoplasm occurs in spherical vesicles-like structures in acapsular *cap67*Δ and melanized H99 cells. Scale, 5 μm.

### Visualizing the reducing-end probes distribution in C. neoformans

Incubation of the reducing-end probe **6** with *C. neoformans* strain H99 in rich or minimal media revealed that the probe localized near the cell membrane – cell wall interface. Incubations with R.E probe **6** and directly conjugated anti-glucuronoxylomannan (GXM) antibody 18B7 (18B7-AF_594_), revealed no probe labelling throughout the capsular GXM (Figure 2C). In melanized fungal cells, we observed stronger fluorescence from the R.E probe in organelles or vesicular bodies around the periphery of large vacuole inside the cells (Figure 2D). Acapsular mutant *CAP67*Δ showed no staining by 18B7-AF_594_, as expected from the absence of capsular GXM, but did manifest fluorescence in the cytoplasm and at the cell wall-membrane interface after labelling with the R.E probe (Figure 2E). These structures could reflect the cytoplasmic accumulation of polysaccharide containing vesicles associated with the acapsular phenotype (38). The presence of R.E probe labelling inside the cytoplasm of the melanized and acapsular mutants suggests the probe can penetrate the cell wall and membrane and react with reducing sugars in the cytoplasmic space. This phenomenon was less evident in encapsulated H99 cells (Figure 2C). It is possible that in melanized cells, the strong cell wall fluorescence observed in non-melanized cells is quenched by the pigment, thus allowing better visualization of cytoplasmic fluorescence and revealing internal details. In addition, we noted that the cell wall of melanized cells contained localized fluorescent signals (Figure 2D) which could reflect vesicles crossing the cell wall, which have been shown to carry polysaccharide (39).

### Incubating Cryptococcus spp. with the reducing-end probe

To examine the capsule architecture across the *Cryptococcus* genus, we incubated two *Cryptococcus gattii* strains representing serotype B (ATCC 24065) (Figure 3D) and C (ATCC 32608) (Figure 3C), an environmental strain *Cryptococcus albidus* (Figure 3B), which has been characterized as having a capsule akin to a serotype A strain (40), and a serotype D strain *C. neoformans* (ATCC 24067)(Figure 3A). While well documented phenotypic differences were observed (41), including differences in capsule diameter, globose, oblong and elliptical cell shapes, all cryptococcal spp. analyzed reacted to the R.E probe in a conserved manner, with the R.E probe localizing at the capsule-wall interface (Figure 3). Furthermore, the incubation of the R.E probe with the various strains tested resulted in similar level of fluorescence at the emission maxima of the probe (Figure 3E).

**Figure 3.**
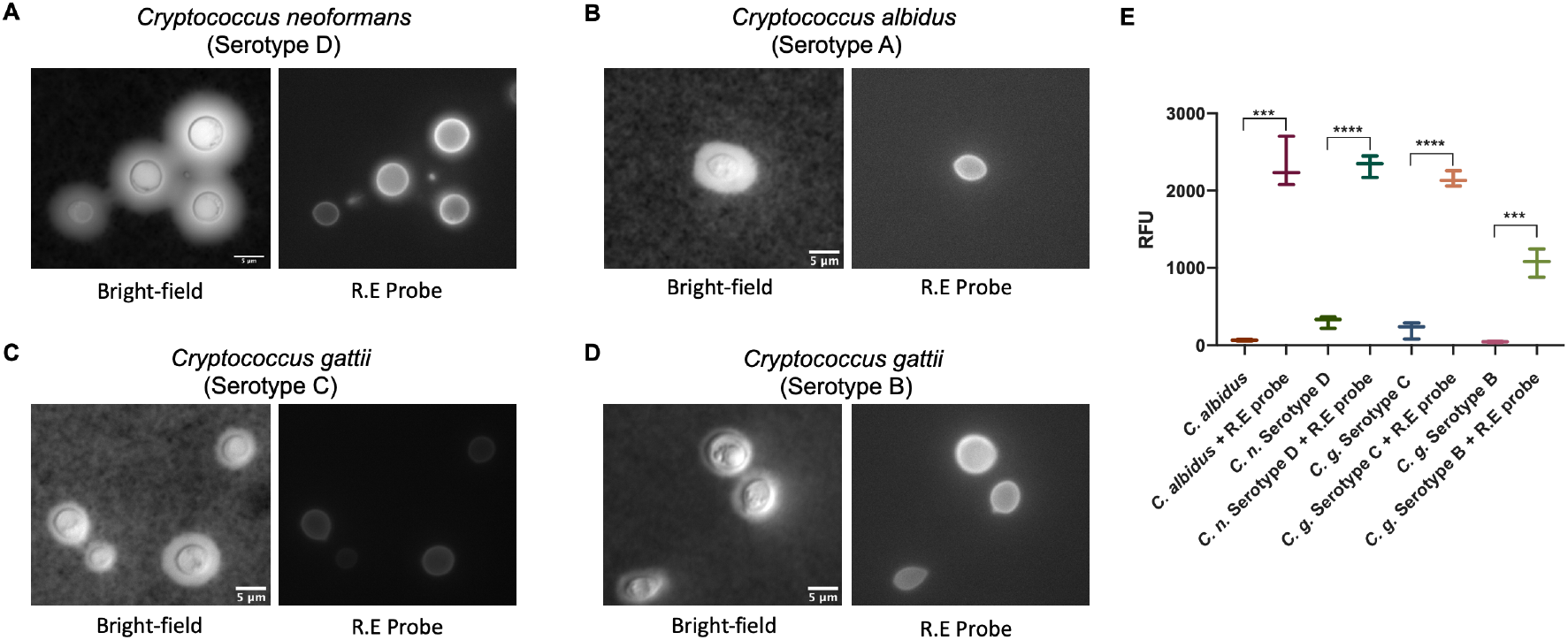
Localization of reducing end polysaccharides is maintained across the *Cryptococcal* genus as label by the R.E probe. Incubations with the R.E probe with any of the *Cryptococcal* spp. tested result in a similar staining pattern. Suggesting that capsule architecture and biosynthesis is maintained across species. **A** *C. neoformans* (ATCC 24067). **B** *Cryptococcus albidus* **C** *Cryptococcus gattii* (ATCC 32608) **D** *Cryptococcus gattii* (ATCC 24065). **E** Incubation with the R.E probe increases RFU at emission of probe (excitation 360 nm), Error bars represent 95% confidence interval, Statistical significance was determined using unpaired T-test (*, *p* 0.05; **, *p* 0.01; ****, *p* 0.0001.). Scale 5 μm.

### Capsule perturbation alters probe reactivity

To better understand the localization of the R.E probe **6**, 3-day-old H99 encapsulated cells grown in minimal media were processed in three ways: DMSO, sonication, and French press. DMSO is used to remove the capsule from the cell (42). French press passage has been used to isolate cryptococcal cell walls from capsular polysaccharide (43). Sonication was previously shown to strip the capsule (44). We hypothesized that these treatments may perturb the capsule structure, which could affect the R.E probe labelling and provide information on selectivity. We also hypothesized that some of the cellular processing methods might expose GXM reducing ends that could then be labelled with the R.E probe **6** (Figure 4). DMSO-treated H99 cells lost the large majority of their GXM capsular polysaccharide, as evident by a diminished signal from the 18B7-AF_568_ conjugate (Figure 4). Treating *C. neoformans* cells with DMSO altered the localization of the R.E probe compared to that of the control. The R.E probe signal became stronger inside the cell, suggesting the cell membrane had been permeabilized by the DMSO treatment, and this affected the permeability and reactivity. The Uvitex-2b fluorescence signal was less sharp compared to that of the control, suggesting the cell wall was also affected in DMSO-treated cells. Importantly both dyes altered in a manner independent from each other (Figure 4), indicating labelling of chitin or chitosan was not the predominant localization of the R.E probe.

**Figure 4.**
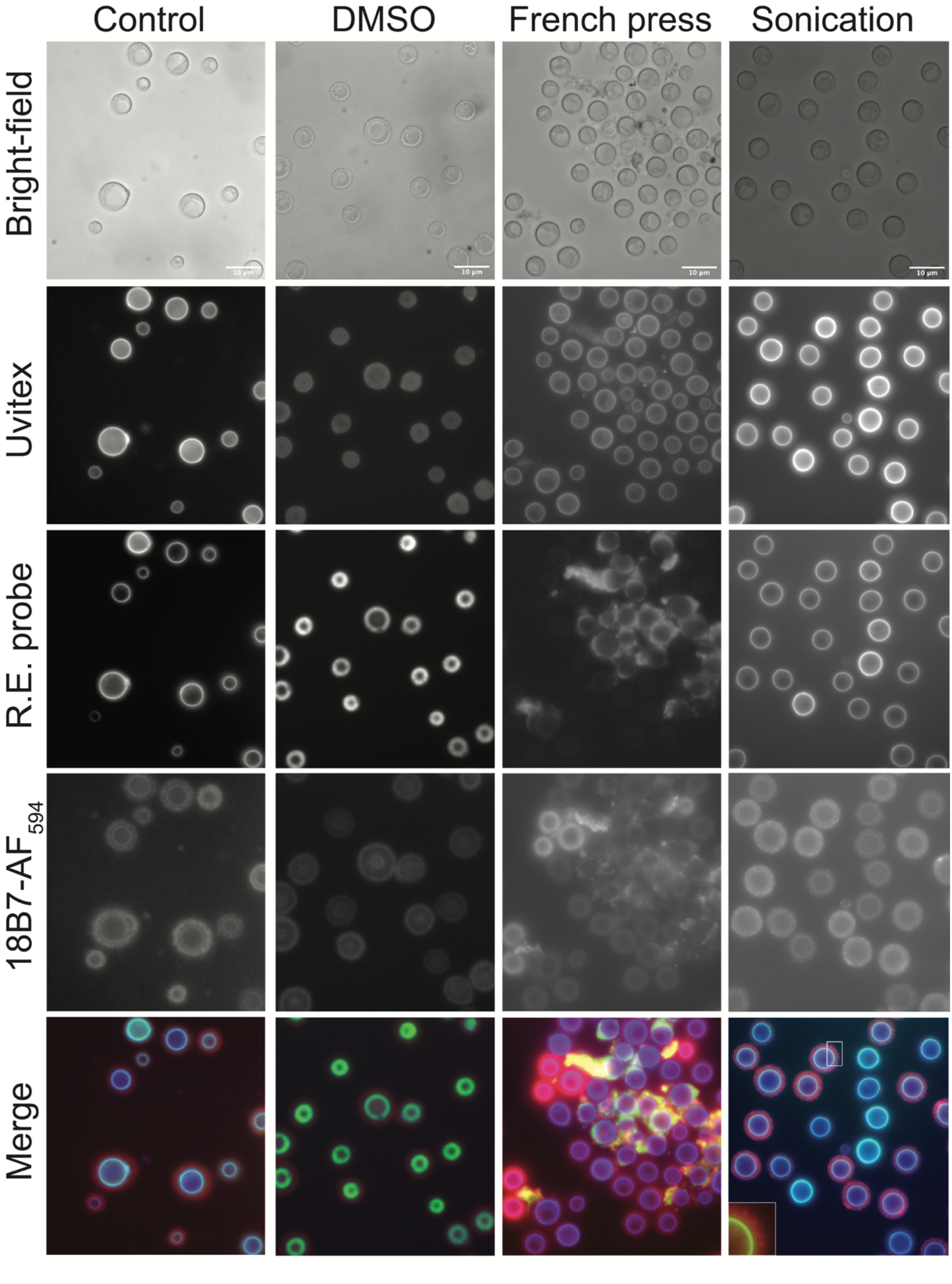
Capsule perturbation alters probe reactivity. H99 *C. neoformans* cells were grown in capsule inducing minimal media for 3 days. Subsquently the cells were independently processed with three different methods: DMSO, French press and sonication. Thereafter the cells were incubated with the flurorecent R.E probe (green) overnight, washed X3, stained for 30 min with Uvitex-2b (Blue) and an 18B7-Alexa Fluor 594 conjugate (red) for 1h. Scale, 10 μm.

Incubation of the R.E probe with French press-treated cells caused the greatest signal disruption of the R.E probe, suggesting the greatest quantities of reducing polysaccharide were exposed by this method. This can be visualized by clusters of sheared capsule polysaccharides that reacted with R.E probe and were not associated with any cell. We interpret this result as implying that the mechanical distribution caused by French press treatment exposed the reducing ends of polysaccharides, that were once localized only at the cell wall, or that French press treatment can cause the breakage of glycosidic bonds due to high temperature and pressures (43). The co-localization signal from 18B7-AF_568_ and the R.E probe showed that the R.E linker labelled the GXM reducing ends (Figure 4). In contrast, the Uvitex-2b signal was not affected significantly by French press treatment (Figure 4). Treating cells with a horn sonicator for 30 seconds (20 Watts) caused the shearing of the capsule, as visualized by the smaller diameter of the capsule compared to the control cells. The localization of the R.E probe was unaffected by this cell-treatment strategy as was the Uvitex-2b binding. However, sonication also reduced a zone of clearance typically observed between the cell-wall and the 18B7-AF_568_ conjugate (Figure 4, merged Sonication, inset)(44). Furthermore, in sonicated cells the R.E probe signal colocalized with 18B7-AF_568_ in a yellow band around the cell body (Figure 4, merged Sonication, inset).

### Extracellular vesicular bodies in CAP59Δ and CAP67Δ mutants

The ability of the reducing-end probe to permeate into the cell and label what appeared to be glycan containing vesicular bodies prompted further investigation. Using two lipophilic dyes BODIPY TR methyl ester (BODIPY TR) and BODIPY FL C_12_ (BODIPY C_12_), we visualized the presence of structures consistent with extracellular vesicular structures or organelles in both *CAP59*Δ and *CAP67*Δ mutants (Figure 5). Furthermore, we visualized extracellular glycan containing vesicles in both *CAP59*Δ and *CAP67*Δ mutants as the R.E probe and lipid dye colocalized extracellularly. (Figure 5A, 5B). Although these acapsular strains do not secrete GXM they do release GalXM (45). We also visualized extracellular structures that stained with BODIPY TR but not with the R.E probe, which presumably represent non-glycan containing vesicles (Figure 5C). Intracellular staining was difficult to resolve as either lipophilic dye caused substantial staining, possibly labelling mitochondria, Golgi apparatus, and vesicular bodies. Despite this the R.E probe did appear to colocalize with the lipophilic dyes, possibly in vesicles or Golgi apparatus. It was also apparent that the R.E probe did not enter the vacuole of the cells or that it was quenched in that environment.

**Figure 5.**
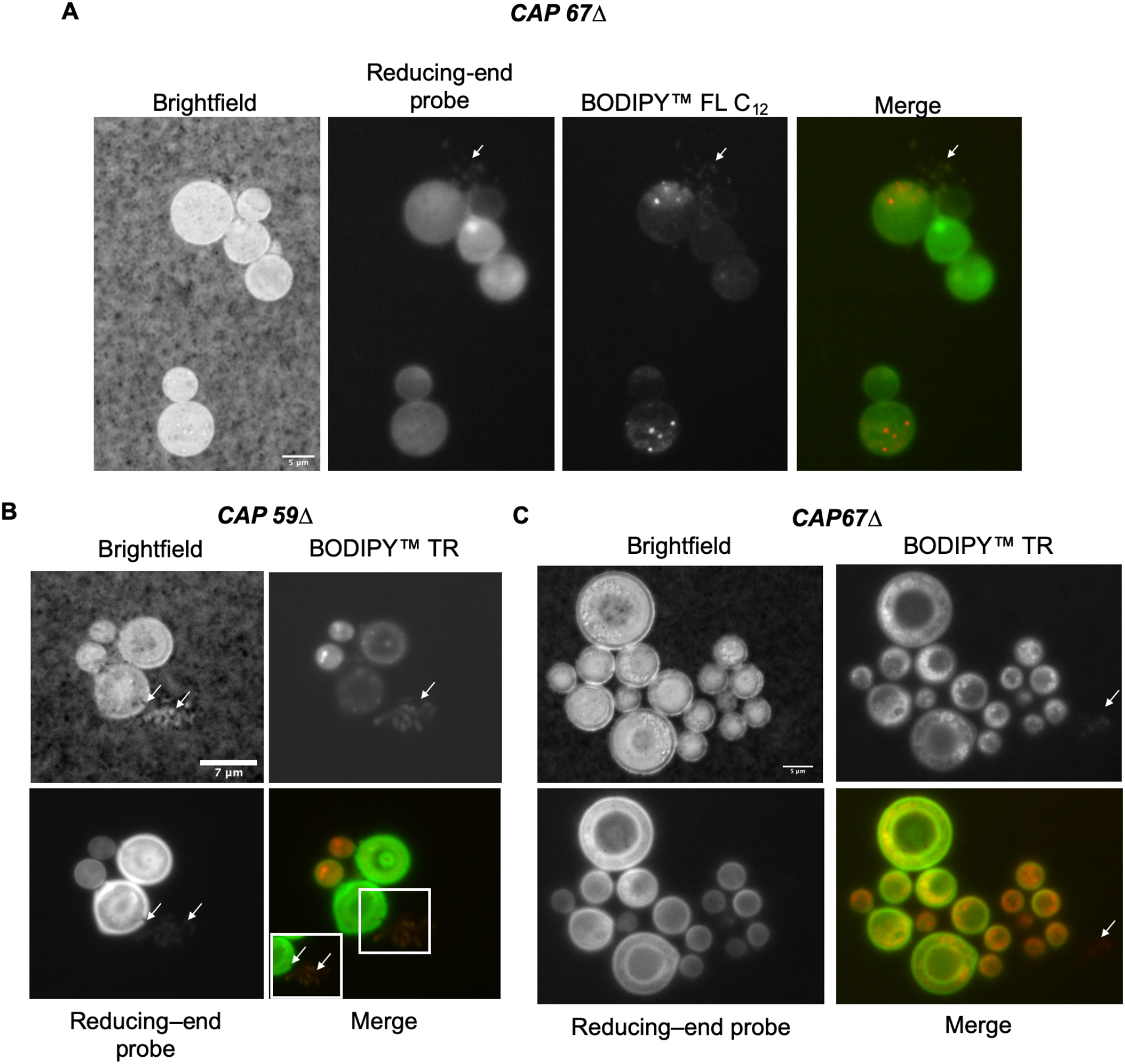
Secretion of vesicular bodies in CAP59Δ and CAP67Δ mutants. **A** Incubation of *CAP67*Δ with BODIPY FL C_12_ and the R.E probe shows vesicle secretion (arrows). **B** *CAP59*Δ cells incubated with BODIPY TR also show signs of secretion of glycan containing vesicles (inset and arrows). **C** *CAP67*Δ incubated with BODIPY TR and R.E probe show signs of colocalization, however BODIPY TR also stains non-glycan containing vesicles (arrows) and shows signs of secretion of non-glycan containing vesicular bodies (arrow). Scale denoted in brightfield: **A** 5 μm, **B** 7 μm, **C** 5μm.

### Dynamic reorganization of Cryptococcus neoformans polysaccharide capsule and cell cell-wall to facilitate budding

Previously, observations of gentle pressure to India-ink stained *C. neoformans* cells caused perpendicular lines to the budding axis suggesting a complex organization of the capsule (46). Hence, we wondered whether we would gain further information around capsular organization using the R.E probe. We subjected encapsulated H99 cells grown in minimal media (M.M) for 2 days to gentle pressure after staining with the R.E probe, 18B7 and Uvitex-2b (Figure 6), to simultaneously stain for reducing end glycans, capsule and cell wall, respectively. We observed the same perpendicular India-ink lines to the budding axis as reported previously (46) but also observed that the capsule, cell wall, and R.E probe had become polarized along the budding axis. This is suggestive that the innermost layer of GXM is attached to the cell wall in a very strong manner, possibly through covalent bonds to either the chitin or glucans. The density of capsule is altered as a result of budding, that allows India-ink to penetrate deeper in perpendicular lines to that of the budding axis. The Uvitex-2b staining correlates to this change in capsule density as the chitin polarizes along the budding axis.

**Figure 6.**
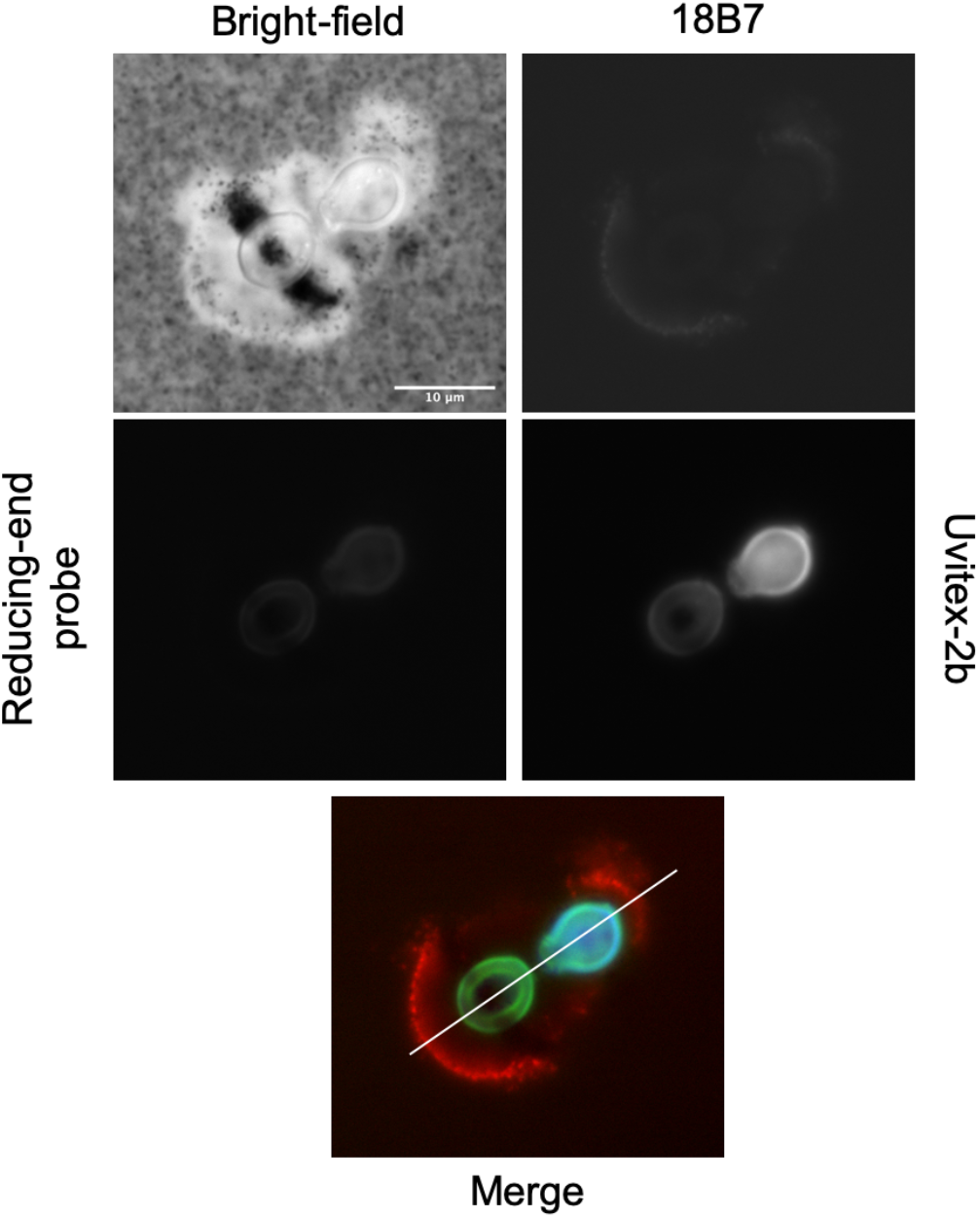
Dynamic reorganization of Cryptococcus neoformans polysaccharide capsule and cell cell-wall to facilitate budding. India ink staining perpendicular to emerging bud correlates to reorganization of chitin cell wall and capsule. 18B7 and R.E probe both have been polarized along the budding axis (line merge). This causes a change in capsule density perpendicular to budding axis allowing India-ink staining to penetrate deeper when gentle pressure is applied to the coverslip. Scale 10 μm.

### Applying shear pressure to understand probes localization

To examine the localization of the R.E probe further, we applied shear pressure to the cover glass slip of encapsulated H99 cells previously labelled with 18B7, Uvitex-2b and the R.E probe. The shear pressure caused some cells to burst or fragment (Figure 7, arrows), but also appear to remove capsular material from cells. This experiment further confirmed that the reducing end probe localized at the base of the GXM polymer at the capsule – cell wall interface (Figure 7). This implies that the capsule in encapsulated cells is organized in a more stable manner than a simple collection of self-aggregated GXM polymers and with strong attachment to cell wall chitin or glucans either through direct linkage to the polysaccharide or proteins that embedded in the cell wall (13, 47).

**Figure 7.**
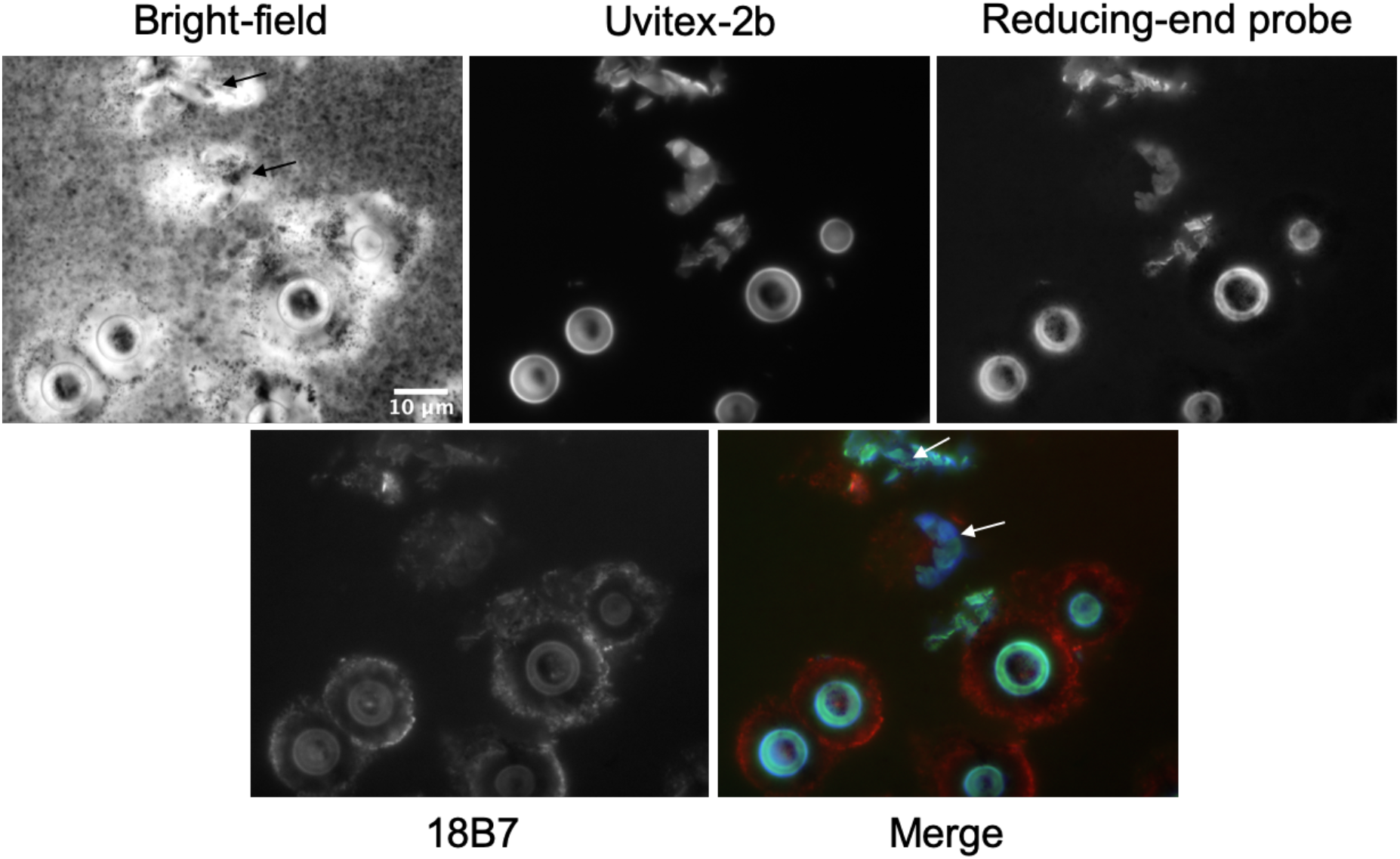
Applying shear pressure to understand probes localization. Application of shear pressure to glass slides caused a population of cells to rupture. Several fragments of cells can be seen (arrows), where capsule can be seen attached to the cell wall, also where the R.E probe localization occurs. Scale 10μm.

### Incubation of EPS and CPS fractions with the reducing-end probe

To further understand the location and abundance of reducing ends in the capsule of *C. neoformans* and its EPS and CPS (Figure 8A), we prepared three fractions for investigation and a chitin sample as a positive control. Corresponding to a 10 kDa, 100 kDa of the exopolysaccharide portion isolated through ultrafiltration membrane discs and a CPS fraction that was prepared by DMSO treatment of the *C. neoformans* cell pellet. Each fraction was then incubated with the R.E probe overnight at 30 °C in the dark and subsequently dialyzed (MWCO 3.5 kDa) to remove the unreacted probe. Aliquots from each sample were then compared to controls to see if there was an increase in fluorescence post-incubation, corresponding to the linker reacting with the reducing end of the sample. All samples that were incubated with the reducing-end linker showed at least a >2-fold increase in RFU compared to controls, consistent with the notion that the linker reacts with reducing sugars (Figure 8B) and forms an oxime conjugate. This experiment also revealed that the EPS fractions shed from *C. neoformans* into the extracellular space contain reducing ends, something not previously known.

**Figure 8.**
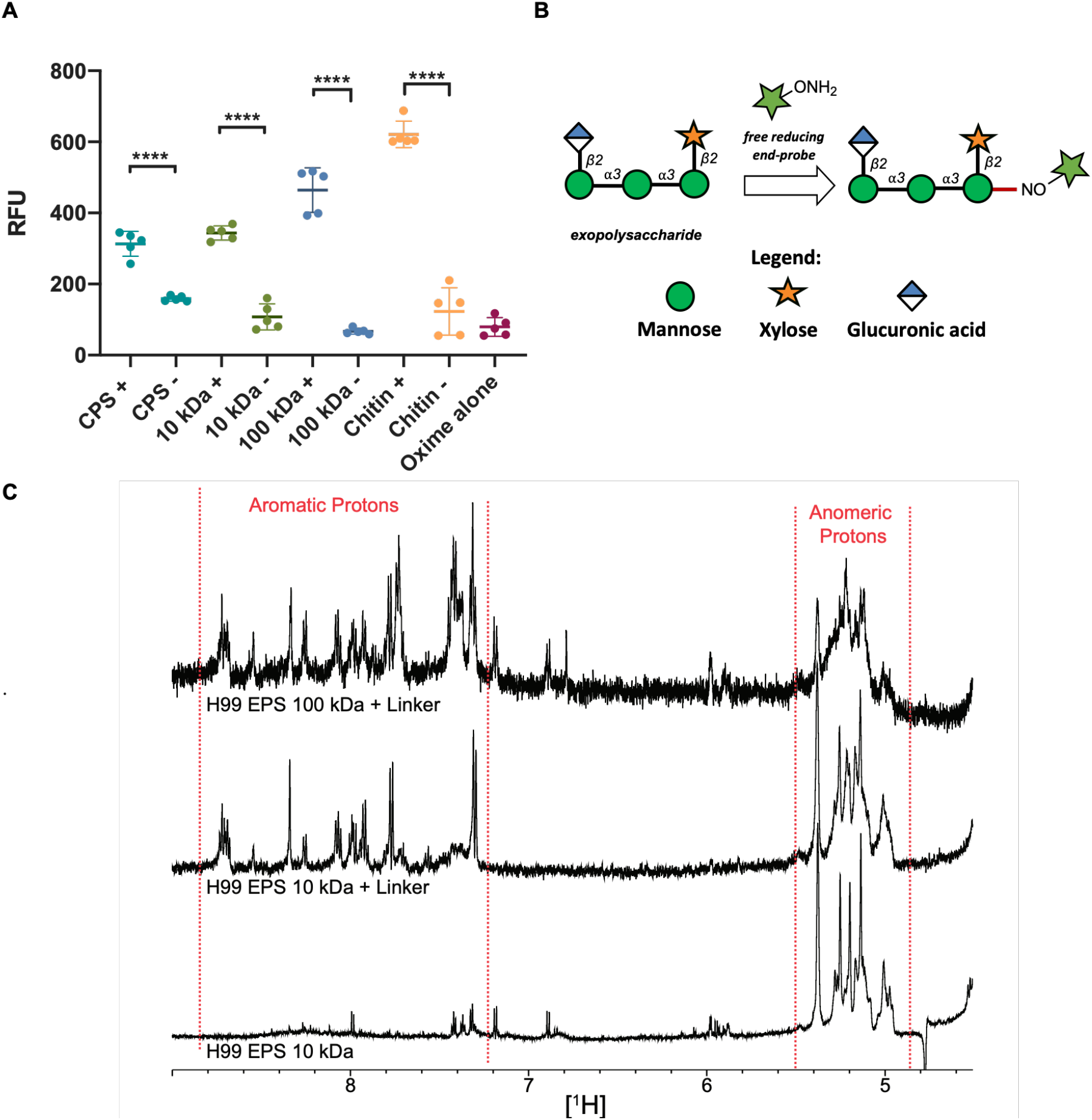
Exopolysaccharides of C. neoformans contain reducing ends. **A** The R.E probe was incubated with CPS, EPS (10 kDa, 100 kDa), Chitin (positive control) or alone for 24 hours. The incubation was repeated in three independent experiments. Following dialysis, fluorescence emission spectra were recorded (excitation 360 nm) and compared to their non-incubated controls, showing an increase in fluorescence at the probe’s emission maxima. Indicating the R.E probe is forming a conjugate with the reducing end of various *C. neoformans* EPS and CPS isolates. Polysaccharides that were incubated with the R.E probe show >2-fold increase in RFU compared to controls, revealing that the shed exopolysaccharide contains reducing ends. Error bars represent 95% confidence interval, Statistical significance was determined using unpaired T-test (*, p 0.05; **, p 0.01; ****, p 0.0001.) B Hypothesis: if reducing ends are present in the exopolysaccharide of C. neoformans they would form stable conjugates with the R.E probe, which could be determined using a fluorescence spectrometer. Glycan notification followed the Symbol Nomenclature for Glycans (SNFG). **C** ^1^H NMR analysis of 10 kDa and 100 kDa exopolysaccharide isolates revealed the presence of aromatic peaks in the region of δ 7.7-8.6 ppm, that are not present in the original EPS fraction. The Exploring *C. neoformans* with a hydroxylamine-armed probe 28 aromatic signals are characteristic of the R.E probe. Characteristic structural reporter groups of anomeric mannose protons can be visualized from δ 5.5-4.9 ppm.

To further establish that the incubated EPS samples did indeed react with the linker, we used ^1^H NMR analysis to confirm the presence of the probe’s aromatic peaks at δ 7.7-8.6 ppm in the spectra of the 10 kDa and 100 kDa EPS fractions (Figure 8C). However, we were unable to obtain an ^1^H NMR of the CPS and chitin carbohydrate signal when samples were run in deuterium oxide, this is likely due to low solubility of chitin in an aqueous solvent and the heterogeneity of the CPS sample (Supporting Information).

## Discussion

We describe the synthesis of a hydroxylamine-armed fluorescent probe that selectively labels reducing sugars, and the use of this probe to study the capsule architecture and biosynthesis in *C. neoformans.* The series of reactions performed yielded significant insights into capsular architecture, biosynthesis, and confirmed that this probe has great potential to help answer outstanding questions in the field. One striking result was the lack of reactivity of the R.E probe with the body of the polysaccharide capsule. Instead, the probe revealed that reducing end glycans are mostly localized at the cell wall-capsule interface. This suggests that these groups are involved in capsule attachment to the cellular surface and/or serve as sites for the origin of capsular assembly. This result would be consistent with a model of capsular assembly occurring at the cell wall and extending away radially. A precedent for this type of capsular biosynthesis and assembly can be found in both eukaryotic and prokaryotic cells. For example, in *Escherichia coli* there are over 80 capsular serotypes, but despite this diversity there is little variation in capsular biosynthesis or assembly (20). In gram-negative bacteria, it is proposed that multiprotein complexes are used to span the cell envelope for the transportation of the capsule to the cell surface (48, 49). In this work, we revealed that despite species differences the R.E probe did not react with the capsule body, suggesting conservation of capsule assembly and biosynthesis processes conserved across *Cryptococcus* spp.

The most striking observation was the absence of R.E. probe reactivity in the body of the capsule. The most straightforward interpretation of this observation is that there are no reducing ends in the body of capsule. However, certain caveats suggest caution in making inferences from this negative result until confirmed by independent approaches. First, GXM macromolecules have mass in the millions of Daltons (50), making reducing end sugars a very small constituent of the molecule and the addition of a single fluorophore may not be detectable relative to the strong signal from the cell wall and cytoplasm. Second, the capsule does contain lipids (51, 52), which can quench fluorescence (53) and it is conceivable that R.E. probe fluorescence is quenched, as may occur in the melanized cell wall. Third, and least probable, would be an accessibility constraint the precluded reactivity. Although GXM macromolecules appear to be oligodendrimers with a very dense core (14), the lack of accessibility explanation is less likely given that the R.E. probe reacted with exopolysaccharide. Mindful of these caveats we proceed with the most parsimonious interpretation of the results and consider the implications of no R.E. probe reactivity in the capsule for its assembly and structure.

An internal location near the cell wall for the synthesis of the capsule was proposed based on the analysis of antibody-labelled capsule polysaccharide movements (54). A subsequent study proposed that capsular enlargement occurred through distal extension although these mechanisms were not mutually exclusive (11). Our observations are consistent with the hypothesis of Frases *et al.* (55) that the distance from the cell body to the capsule is spanned by the length of a single GXM polymer (Figure 9), which is also supported by the finding that capsule size is regulated at the polymer level (56). The orientation of this polysaccharide can be deduced by the reactivity of R.E probe such that the capsule’s reducing end faces into the extracellular space (Figure 9). This observation suggests a model where some capsule biosynthesis may occur at the cell wall, possibly through enzymes that spool the growing polysaccharide chain out into the extracellular milieu. The vesicular polysaccharide containing vesicles may contain the seed motif required for cell wall associated enzymes to material to elongate the structure. An analogous motif to the core pentasaccharide in required in *N*-glycan biosynthesis in humans (57). Taken together with our results that reducing ends occur at the cell wall - capsule interface, this adds evidence supporting the “proximal growth” capsular assembly hypothesis.

**Figure 9.**
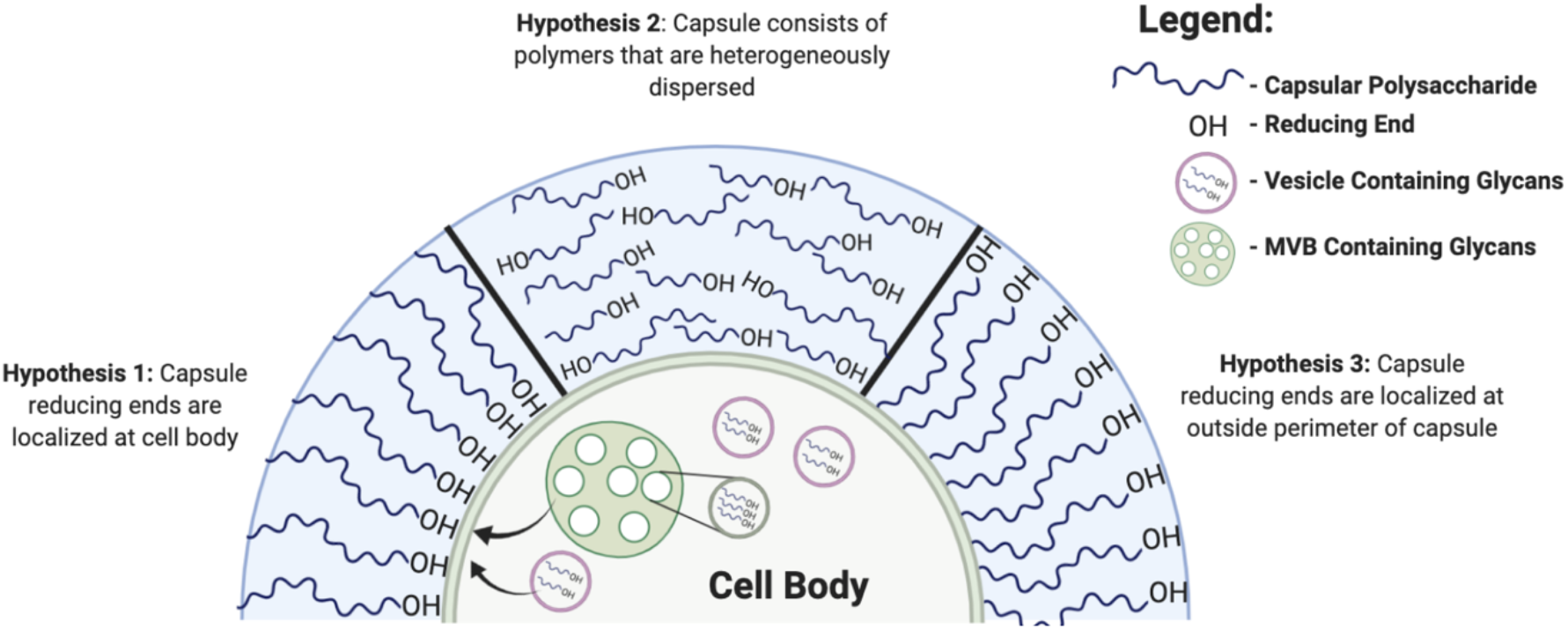
Diagram illustrating three hypotheses for capsule assembly in C. neoformans. The most parsimonious interpretation of the R.E. Probe data supports Hypothesis 1.

The synthesis of capsular polysaccharides was shown to occur within vesicular bodies, and the export of these vesicles across the cell wall is necessary for the export of polysaccharides and capsule assembly (23, 58, 59). Furthermore, extracellular vesicles are implicated in diverse functions, including cell-communication and transport of virulence factors (60, 61). We gained additional evidence of the importance of vesicles in *C. neoformans* capsule biosynthesis in both acapsular mutants *CAP67*Δ, *CAP59*Δ, and melanized H99 cells. *CAP59*Δ and *CAP67*Δ are believed to have defective transport mechanisms for capsule assembly and manifested R.E probe in spherical vesicle structures in the cytoplasm that were not observed in H99 encapsulated cells. We suspect that these structures contained high quantities of reducing glycans, possibly capsular polysaccharides that accumulated in the cytoplasm as a result of the export defect associated with the acapsular phenotype (38). Vesicular structures were also observed inside melanized *C. neoformans* cells around the periphery of a large vacuole. Furthermore, the punctate pattern of fluorescence observed in the melanized cell wall could reflect polysaccharide carrying vesicles across this structure. It is noteworthy that the R.E probe was able to pass through the melanized cell wall, and that it was not sequestered in melanin due to its aromatic and hydrophobic nature. Despite the defective mechanisms for vesicle transport in the *CAP59*Δ and *CAP67*Δ mutants, we observed vesicle secretion in these mutants of both glycan containing vesicles and other vesicles not stained by the R. E. probe. While glycan containing vesicles could represent partial capsular assembly, other vesicles could contain proteins and RNA, which have recently shown to be important in virulence of *Cryptococcus gattii* (61). Intracellular staining with lipophilic dyes was more complex and difficult to resolve, although co-localization did appear, staining of other cellular organelles was present, and due to the presence of large vacuoles in cells, it made total resolution difficult.

Our observations with the R.E probe add to the wealth of data indicating the importance of vesicles in polysaccharide transport and capsule assembly. Further, we believe that the R.E probe could be of great use to delve further into the mechanism of polysaccharide containing vesicle transport and polysaccharide integration into the capsule itself utilizing microscopic techniques.

Further evidence of the capsules highly organized and dynamic structure was revealed by using a technique previously described by Zaragoza *et al* (46). In this work we observed the same ring-like channels observed in the work by Zaragoza and colleagues. Further, the cell wall chitin, reducing ends, and distal capsule (as stained by 18B7) polarize along the budding axis. Together, this suggests a coordinated alteration to both the cell wall and capsular architecture is necessary during the budding process.

Several questions about *C. neoformans* EPS revolve around its function and origin. It has been posited that EPS is the loss of capsular polysaccharide due to rearrangement, or that it is unincorporated capsular polysaccharides (10, 62, 63). While Frases *et al* showed a number of physiochemical differences between EPS and capsular polysaccharides, no report has observed reducing ends in EPS. Incubation of a 100 kDa and 10 kDa exopolysaccharide fraction confirmed the presence of reducing ends in the polysaccharides that are shed into the extracellular space. This has particularly important implications for vaccine development, as it provides a functional group that may be exploited in creating glycoconjugates for vaccine development. Furthermore, the presence of a reducing group in soluble exopolysaccharide could greatly facilitate their attachment to surfaces such as beads for immunological assays and purification of antibodies using affinity chromatography.

We also gained insight into how different cellular processing methods affect the polysaccharide capsule as visualized through microscopy. Overall, processing the *C. neoformans* cells with DMSO, French press, and sonication resulted in different effects on capsule size and membrane-wall integrity as measured by R.E probe reactivity. DMSO treated cells had absent or reduced capsules, and there were indications that the cell membrane was permeabilized. Similarly, sonicated cells displayed reduced capsule size, but in contrast, membrane structure had not been affected. In the sonicated cells, it was also possible to visualize the co-localization of the R.E probe and the 18B7_AF568_, conjugate in the capsular polysaccharide. French press exposed reducing ends of the capsule that were previously associated with the cell wall or inaccessible and were then labelled by the R.E probe and 18B7_AF568_, providing further confirmation that the probe can react with GXM.

Given that reducing ends (aldehydes) are among the most reactive groups in polysaccharides, the lack of R.E probe labeling in the capsule suggests that this could be a strategy to reduce capsular reactivity to compounds in soil and/or phagosomes after ingestion by phagocytic predators such as amoeba. For example, polysaccharide-reducing groups can react with protein amino groups in the Maillard reaction, which could potentially generate toxic compounds or produce stresses that damage the capsule. Microbial polysaccharides are vulnerable to depolymerization by reactive oxygen and nitrogen species, and the presence of reactive reducing groups enhances their susceptibility to a myriad of microbicidal compounds produced by the oxidative burst of neutrophils, macrophage, and amoeba (64).

In summary, our results establish a proof of concept that chemical biology provides new tools to study the capsule of *C. neoformans*. For example, the R.E probe could be used to label extracellular vesicles containing polysaccharide and thus facilitate their purification. In further studies, we intend to identify and characterize the glycan structures containing reducing ends and hope to possibly use insights for the development of novel therapeutics against *C. neoformans.*

## Experimental procedures

### General synthetic methods

Unless otherwise noted all reactions containing air- and moisture-sensitive reagents were carried out under an inert atmosphere of nitrogen in oven-dried glassware with magnetic stirring. N_2_-flushed stainless cannulas or plastic syringes were used to transfer air- and moisture-sensitive reagents. All reactions were monitored by thin-layer chromatography (TLC) on Merck DC-Alufolien plates precoated with silica gel 60 F254. Visualization was performed with UV-light (254 nm) fluorescence quenching. Evaporation in vacuo/under vacuum refers to the removal at 40 °C, unless otherwise stated, of volatiles on a Buchi rotary evaporator with an integrated vacuum pump.

### Chromatography

Silica gel flash chromatography was carried out using Davisil LC60A (40-63 μm) silica gel or with automated flash chromatography systems, Buchi Reveleris® X2 (UV 200-500 nm and ELSD detection, Reveleris® silica cartridges 40 μm, BÜCHI Labortechnik AG) and Biotage® SP4 HPFC (UV 200-500 nm, Biotage® SNAP KP-Sil 50 μm irregular silica, Biotage AB).

### Synthetic Materials

All chemicals for the synthesis were purchased from commercial suppliers (Acros, Carbosynth Ltd, Fisher Scientific Ltd, Glycom A/S, Merck, Sigma-Aldrich, VWR) and used without purification. Dry DCM and THF were obtained from a PureSolv-ENTM solvent purification system (Innovative Technology Inc.). All other anhydrous solvents were used as purchased from Sigma-Aldrich in AcroSeal® bottles.

First, inclusion of the probe TCI triple gradient, and the internal standard D6DSS at 0. I also tend to include the pulse programs used,

### Instrumentation

^1^H NMR (400, 500 or 600 MHz), 13C NMR (101 MHZ or 125 MHz) spectra were recorded on Varian-inova or Bruker spectrometers at 25 °C in chloroform-d1 (CDCl_3_), methanol-d4 (CD_3_OD), water-d2 (D_2_O), ^1^H NMR spectra were standardized against the residual solvent peak (CDCl_3_, δ = 7.26 ppm; CD_3_OD, δ = 3.31 ppm; D_2_O, δ = 4.79 ppm; d6-DSS δ = 0.0 ppm or internal tetramethylsilane, δ = 0.00 ppm). Bruker instrumentation spectrometers were equipped with Avance II console and triple resonance, TCI cryogenic probe with z-axis pulsed field. ^13^C NMR spectra were standardized against the residual solvent peak (CDCl_3_, δ = 77.16 ppm; CD_3_OD, δ = 49.00 ppm. All ^13^C NMR are ^1^H decoupled. All NMR data is represented as follows: chemical shift (δ ppm), multiplicity (s = singlet, d = doublet, t = triplet, q = quartet, dd = doublet of doublets, ddd = doublet of doublets of doublets, dt = doublet of triplets, m = multiplet, br = broad signal, ad = apparent doublet, at = apparent triplet), coupling constant in Hz, integration. Assignments were aided by homonuclear ^1^H−^1^H (COSY, TOCSY), and ^1^H-^13^C heteronuclear (HSQC, HMBC) two-dimensional correlation spectroscopies. ^13^C chemical shifts were reported with one digit after the decimal point, unless an additional digit was reported to distinguish overlapping peaks. Software for data processing: MestReNova, version 11.0.0-17609 (MestReLab Research S.L.). High-resolution mass spectrometry (HRMS) data were recorded on a Waters micromass LCT LC-Tof instrument using electrospray ionization (ESI) in either positive or negative mode. Low-resolution mass spectrometry (LRMS) experiments were recorded on a Waters micromass Quattro Micro LC-MS/MS instrument using electrospray ionization (ESI) in either positive or negative mode.

### Specific details for a synthetic procedure can be found in the supporting information

#### Growth conditions

*C. neoformans* Serotype A strain H99 (American Type Culture Collection (ATCC) 208821), *C. neoformans* Serotype D (ATCC 24067), *Cryptococcus gattii* strains serotype B (ATCC 24065) and C (ATCC 32608), *CAP59*Δ, *CAP67*Δ, *Cryptococcus albidus* was grown for 48 h at 30 °C in capsule inducing media composed of: 10 mM MgSO_4_, 29.3 mM KH_2_PO_4_, 13 mM glycine, 3 μM thiamine-HCl; adjusted to pH 5.5 and supplemented with 15 mM (regular minimal media) dextrose.

#### Cellular processing techniques

DMSO extraction carried out as previously described (65). Ultrasonication carried out as previously described (44). Isolation of EPS samples (10 kDa, 100 kDa) was carried out as described (13). French press was carried out as previously reported (43)

#### Hydroxylamine-armed probes physicochemical properties incubation with C. neoformans

*C. neoformans* cells were grown inoculated in Sabouraud dextrose medium for 2 days and then transferred to capsule inducing minimal media for 3 days. Cells were pelleted (2000 rpm, 5 minutes) and washed with PBS x3. The cells were then incubated with the reducing end probe overnight, in the dark at 30 °C. Cells were pelleted and washed to the remove excess probe. The spectrometric analysis was completed using a SpectraMax M5 microplate reader. Statistical analysis was performed in GraphPad Prism 8.

#### Fluorescent microscopy

H99, *CAP59*Δ or *CAP67*Δ *C. neoformans* cells were grown inoculated in Sabouraud dextrose medium for 2 days and then transferred to capsule inducing minimal media for 3 days. Samples were blocked in blocking buffer (1% bovine serum albumin (BSA). Thereafter the cells were incubated with or without the 2.5 μM of the fluorescent reducing-end probe overnight at 37 °C in the dark. If desired, a 1:1000 (1 mM) dilution of BODIPY™ FL C_12_ was incubated 5 hours after the addition of the reducing end probe and left overnight. Cells were pelleted and washed to remove excess probe or dye, X3 PBS plus 1% BSA. Stained for 20 min with or without Uvitex-2b, washed X3, with PBS. If desired a 1 mM (1:1000) CellTrace™ BODIPY™ TR Methyl Ester was added 1h prior to imaging. Then incubated with an 1 μg/mL IgG1 18B7-Alexa Fluor 594 conjugate for 1 h, at 37 °C. Channel exposure: FITC (800 ms) (ex:em 498/516 nm), TRITC (600 ms) (ex:em 540/580 nm) and DAPI (50 ms) (ex:em 350/450 nm). Images were collected with an Olympus AX70 microscope, photographed with a QImaging Retiga 1300 digital camera using the QCapture Suite V2.46 software (QImaging, Burnaby BC, Canada), and processed with ImageJ or Fiji (NIH, USA).

#### Polysaccharide labelling and fluorescence reading

CPS, 100 kDa EPS, 10 kDa EPS and Chitin fractions (~1.18 μmoles, 1 eq) were incubated with hydroxylamine-armed probe 1 (1 mg, 2.36 for μmoles, 2 eq) in 1 mL PBS (100 mM, pH 4) for 24 h, shaking, at 30 °C in the dark (set up in triplicate). The reaction mixtures were then transferred to dialysis tubing (1-kDa or 3.5-kDa MWCO, Spectrum laboratories inc., USA) and dialyzed against distilled water for 48 h at room temperature, in the dark, with water replacement every 6 h. Three aliquots of each fraction were then transferred to an opaque plate reader to analyze fluorescence. Fluorescence in different fractions was measured with the SpectraMax M5 microplate reader (Molecular Devices). All measurements were performed at 37 °C with an excitation wavelength of 360 nm, an emission wavelength of 440 nm, and a cut-off emission filter of 435 nm. Wells were set up in triplicate, and 30 readings were taken per well with a 5-s mix time prior to reading. Photomultiplier tube sensitivity was set to high. Statistical analysis was performed in GraphPad Prism 8.

#### Generating Indian ink equatorial rings in Cryptococcus neoformans polysaccharide

Cells were prepared as described above. 2 μL India-ink was added to 8 μL of solution of cells on a glass slide. The coverslip was placed on top and a firm pressure was applied.

#### Rupture of Cryptococcus neoformans cells

Cells were prepared as described above, with Indian-ink stain. Shear pressure was applied by moving the coverslip back and forth four times.

## Acknowledgements

We thank Dr Yannick Ortin for NMR support, Daniel Quinn Smith for the contribution of editing the figures and the Johns Hopkins Biomolecular NMR Center.

## FOOTNOTES

C.J.C was funded by Irish Research Council postgraduate award (GOIPG/2016/998). R.J.B.C was supported by the JHU CFAR NIH/NIAID fund P30AI094189. S.O was supported by Science Foundation Ireland Award 13/IA/1959. A.C. was supported by grant 5R01HL059842.

The abbreviations used are:

R.E probe: reducing-end probe
hydroxylamine: armed fluorescent probe
CPS: capsular polysaccaride
EPS: exopolysaccaride

